# Sodium-Calcium Exchanger Mediates Sensory-Evoked Glial Calcium Transients in the Developing Retinotectal System

**DOI:** 10.1101/2020.11.14.383125

**Authors:** Nicholas J. Benfey, Vanessa J. Li, Anne Schohl, Edward S. Ruthazer

## Abstract

Various types of sensory stimuli have been shown to induce calcium elevations in glia. However, a mechanistic understanding of the signalling pathways mediating sensory-evoked activity in glia in intact animals is still emerging. Here we demonstrate that during early development of the Xenopus laevis visual system, radial astrocytes in the optic tectum are highly responsive to sensory stimulation. Calcium transients occur spontaneously in radial astrocytes at rest and are abolished by silencing neuronal activity with tetrodotoxin. Visual stimulation drives temporally correlated increases in the activity patterns of neighbouring radial astrocytes. Following blockade of all glutamate receptors, visually-evoked calcium activity in radial astrocytes is enhanced, rather than suppressed, while the additional blockade of either glutamate transporters or sodium-calcium exchangers (NCX) fully prevents visually-evoked responses. Additionally, we demonstrate that blockade of NCX alone is sufficient to prevent visually-evoked responses in radial astrocytes, highlighting a pivotal role for NCX in glia during development.

## Main

Following the initial discovery that astrocytes in culture actively respond to glutamate and neuronal activity through increases in internal calcium concentration^1–3^, there has been an expanding effort to understand glial participation in neural circuits *in vivo*^4^. Once largely accepted to be passive cells acting principally as scaffolds for neuronal migration and mediators of ionic and metabolic homeostasis^5^, glia are now understood to be highly active contributors to neural circuit development^6,7^, function^8,9^, plasticity^10,11^, and behaviour^12^, both in health and disease^13^. However, despite experiments performed in adult animals demonstrating that glia respond to physiological stimulation *in vivo*^14,15^, a comprehensive mechanistic understanding of how glia participate in neurotransmission remains elusive^16^. Additionally, an understanding of how sensory stimulation regulates glial calcium signaling during early development remains largely unexplored in intact animals.

Early work on the physiological regulation of glial calcium signaling *in vivo* identified multiple mechanisms underlying glutamatergic signaling between neurons and glia. In barrel cortex of adult mice, the responsiveness of astrocytes to whisker stimulation was shown to depend on type-1 metabotropic glutamate receptors^14^ (mGluR1 and mGluR5), while in the primary visual cortex of adult ferrets, visual responses in astrocytes were reduced following blockade of glutamate transporters^15^. In the cerebellum of adult mice, the response of Bergmann glia to motor activity can be largely abolished by broad spectrum blockade of glutamate receptors^17^, and the calcium activity in Müller glia during retinal waves in zebrafish has been shown to rely on the activation of calcium-permeable AMPA receptors and glutamate transporters^18^. Recent work has also demonstrated that mGluR5 is developmentally regulated, with expression levels in astrocytes decreasing dramatically into adulthood, thereby calling into question the mechanism by which it influences mature astrocytes^19^.

Recent work investigating the physiological activation of glia *in vivo* shows that glutamate receptors play a limited role in generating glial calcium events thereby suggesting the involvement of additional mechanisms^18,20–25^. It is also unclear whether glial calcium responses result from direct activation of glial receptors or occur indirectly following activation of neuronal receptors and release of diffusible signals, as few experiments have employed simultaneous dual imaging of neurons and glia in their preparations. Recent work both *in vitro* and *ex vivo* has suggested that the rapid entry of sodium via glial glutamate and GABA transporters during electrical or pharmacological stimulation of neural circuits may drive extracellular calcium entry into glia through reversal of sodium-calcium exchangers (NCX), thereby linking neurotransmitter reuptake to glial calcium events through a mechanism independent of glutamate receptors^21,26–31^. We have therefore investigated the contribution of NCX to physiologically evoked glial calcium responses *in vivo^32^*.

The development of genetically-encoded calcium indicators has led to significant advances in *in vivo* calcium imaging^33^. As such, we have produced GCaMP6s-expressing albino *Xenopus laevis* tadpoles which are easily amenable to acute pharmacological manipulation and allow for the simultaneous imaging of calcium activity in both neurons and glia in the retinotectal circuit of intact animals during visual stimulation. To dissect the signaling mechanisms by which visual activity regulates calcium events occurring in both tectal neurons and radial astrocytes during early development, we employed a systematic approach involving pharmacological activation and antagonism of glutamate receptors, glutamate transporters, and NCX in conjunction with the presentation of a behaviourally relevant visual stimulus while imaging the optic tectum of GCaMP6s-expressing tadpoles.

We found that calcium events occur spontaneously in both tectal neurons and radial astrocytes during resting state and are abolished by the voltage-gated sodium channel blocker tetrodotoxin. Presentation of a visual stimulus increased calcium events occurring in both tectal neurons and radial astrocytes with the onset of glial responses occurring several seconds after those in neurons. Pharmacological activation of AMPA receptors or mGluR1, but not mGluR5, induced calcium events in radial astrocytes with a delayed onset of tens of seconds to minutes relative to neighbouring tectal neurons. Visually driven increases in radial astrocyte calcium events were not prevented, but rather enhanced, by broad spectrum blockade of both ionotropic and metabotropic glutamate receptors in the optic tectum, but were eliminated following the blockade of excitatory amino acid transporters (EAAT) 1 and 2 or NCX. Blockade of NCX alone was sufficient to prevent radial astrocytes from responding to visual stimulation, thereby demonstrating a critical role for NCX in mediating sensory driven calcium events in glia during development.

## Results

### Radial astrocytes exhibit spontaneously occurring calcium transients in the optic tectum

Radial astrocytes form the scaffold of the optic tectum in *Xenopus laevis,* having at least one large radial process that extends from the ventricular to the pial surface where it forms a broad endfoot. Radial astrocyte endfeet are easily identifiable structures that tile the surface of the brain. Their columnar processes extend numerous filopodia into the neuropil that contact synapses formed between retinal ganglion cell (RGC) axons and tectal neuron dendrites (Fig.1a,b,c)^34,35^. Thus in the amphibian optic tectum radial astrocytes are believed to serve the roles performed in mammalian brain by mature astrocytes. To generate tadpoles that express GCaMP6s in all the neurons and glia throughout the optic tectum, allowing for simultaneous monitoring of both neuronal and glial calcium activity *in vivo,* we performed microinjection of GCaMP6s mRNA in two-cell stage embryos following *in vitro* fertilization. The dynamic calcium fluctuations occurring in neurons and glia were then imaged in these GCaMP6s-expressing tadpoles at developmental stage 47, when considerable retinotectal circuit remodeling is underway, using *in vivo* resonance scanning two-photon microscopy. We found that radial astrocytes exhibit spontaneously occurring calcium transients in their processes and endfeet (Fig. 1d and Video 1). Calcium signal in the endfoot of a radial astrocyte correlated highly with its neuropil signal (Fig. 1e). Regions of interest representing individual tectal neuron somata and radial astrocyte endfeet were identified using an automated segmentation algorithm and fluorescence traces were analyzed for neurons and glia (Fig. 1f,g and Video 2).

**Fig. 1:**
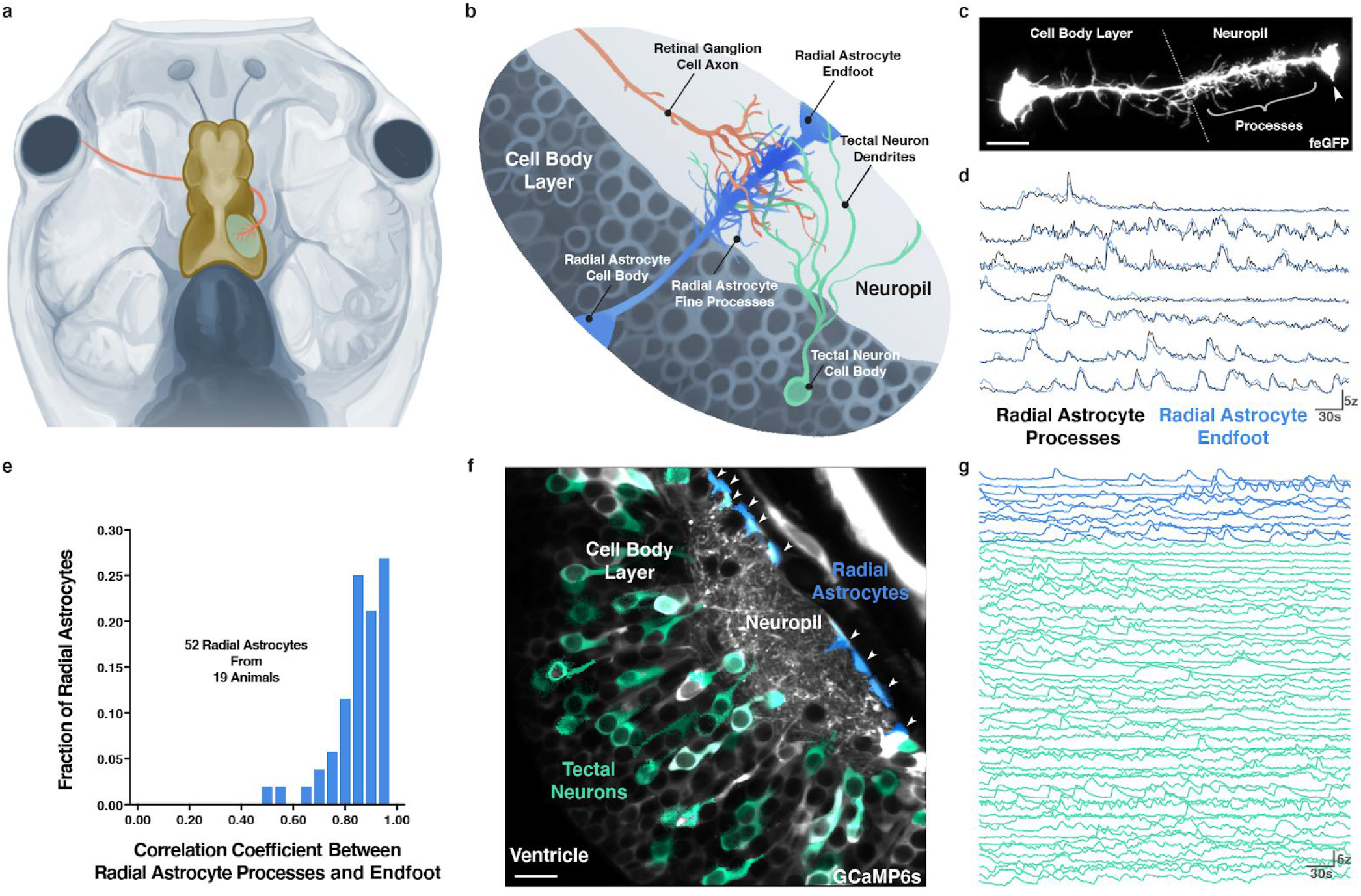
Radial astrocytes in the optic tectum of *Xenopus laevis* exhibit spontaneously occurring Ca^2+^transients during early development. **a**, Schematic of the *Xenopus laevis* tadpole retinotectal circuit showing a retinal ganglion cell axon (red) innervating the contralateral optic tectum (yellow). **b**, Cellular organization of the optic tectum showing the spatial relationships between radial astrocytes (blue), presynaptic retinal ganglion cell axons (red), and postsynaptic tectal neuron dendrites (green). **c**, Average projection of a single radial astrocyte expressing membrane-targeted farnesylated EGFP, arrowhead pointing to the cell’s endfoot (scale bar = 10 μm). **d**, Representative traces of resting state Ca^2+^ activity occurring in radial astrocytes where both their processes (black) and endfeet (blue) are visible in the neuropil, total time = 300 s, z = z-score. **e**, Histogram showing high degree of correlation between radial astrocyte processes and their endfeet. **f**, Averaged temporal projection of 4500 frames (t) from a single optical plane through one hemisphere the optic tectum of a GCaMP6s-expressing tadpole. Regions of interest generated using Suite2p showing active tectal neuron cell bodies (green) and radial astrocyte endfeet (blue, arrowheads) (scale bar = 25 μm). **g**, Representative traces of resting state Ca^2+^ activity from all active cells in a single plane of the optic tectum, radial astrocyte endfeet (blue), tectal neuron cell bodies (green), total time = 300 s. Credit: Artwork in Fig.1a and Fig.1b, A. Desaulniers, Orcéine.

During resting state in darkness, both cell types exhibited extensive calcium activity. We quantified the spontaneous calcium events occurring during 5 min in darkness in 200 radial astrocytes and 390 tectal neurons from 21 animals and found that on average there were 6.00 ± 0.64 (mean ± SEM) active glial endfeet and 18.57 ± 2.02 spontaneously active neurons per imaging plane in each tectum. Over 5 min of imaging, 1.65 ± 1.29 calcium events per glial cell and 2.93 ± 1.18 calcium events per neuron were observed. Together this demonstrates that calcium events occur spontaneously in radial astrocytes during early development and that these events can be monitored in tandem with those in nearby neurons during live imaging *in vivo,* allowing for high temporal and spatial resolution investigation of neuron-glia communication in the developing brain.

### Calcium transients in radial astrocytes are modulated by neural activity

We first sought to determine whether the calcium transients that occur spontaneously in glia during resting state were coupled to neuronal activity (Fig. 2). Tetrodotoxin (TTX) is a potent and selective blocker of the voltage-gated sodium channels that underlie action potential generation in neurons. As voltage-gated sodium channels do not mediate physiological signaling in astrocytes^36^, TTX can specifically silence neuronal spiking. A thin incision through the skin on the head of the tadpole was made to expose the ventricular space and permit rapid pharmacological manipulation of the optic tectum during live imaging (Fig. 2a). After imaging a baseline of calcium activity over a 5-minute period in darkness, either vehicle control (Fig. 2b,d) or TTX (1 μM, Fig. 2c,e) was applied to the brain and a second 5 min period of live imaging was carried out (Fig. 2b-e and Videos 3,4). Application of TTX significantly reduced calcium activity in both neurons and radial astrocytes, as revealed by a decreased number of active cells (Fig. 2f,g) and events (Fig. 2h,i) per animal or per cell (Fig. 2j,k). These results suggest that the calcium transients occurring in radial astrocytes are dependent on neuronal spiking activity.

**Fig. 2:**
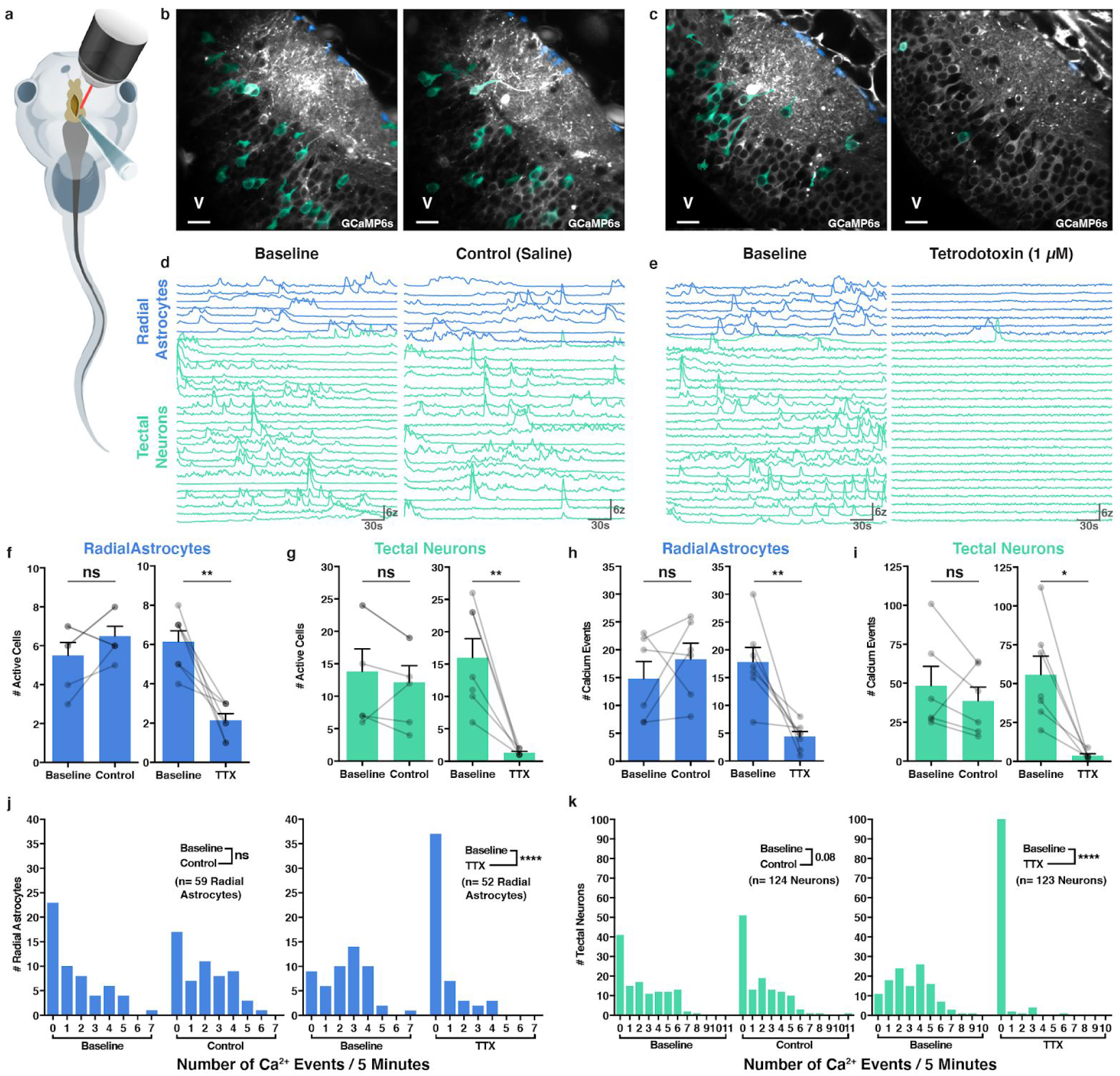
Resting state Ca^2+^ transients in radial astrocytes are coupled to neuronal activity. **a**, Schematic of a *Xenopus laevis* tadpole with a slit cut through the skin into the ventricle to expose the optic tectum to acute pharmacological manipulation during live imaging. **b**, **c**, Average temporal projections, 4500 frames (t), control treatment group (external saline) (**b**), tetrodotoxin treatment group (1 μM) (**c**) (V= ventricle, scale bar = 25 μm). **d**, **e**, Traces of resting state Ca^2+^ activity from control (**d**) and tetrodotoxin (**e**) treated animals, radial astrocytes (blue), neurons (green), total time = 300 s. **f**, **g**, Number of active cells during each imaging period, radial astrocytes (blue), neurons (green) (coloured bars = mean + s.e.m.). Paired t-tests, control, n = 6 animals, radial astrocytes p = 0.2031, neurons p = 0.3632. TTX, n = 7 animals, radial astrocytes **p = 0.0027, neurons **p = 0.0059. **h**, **i**, Total number of Ca^2+^ events during each imaging period, radial astrocytes (blue), neurons (green) (coloured bars = mean + s.e.m.). Paired t-tests, control, n = 6 animals, radial astrocytes p = 0.4121, neurons p = 0.3933. TTX, n = 7 animals, radial astrocytes **p = 0.0065, neurons *p = 0.0125. **j**, **k**, Number of events per cell during each imaging period, radial astrocytes (blue), neurons (green). Mann-Whitney tests, control, n = 59 radial astrocytes and 124 neurons from 6 animals, radial astrocytes p = 0.1876, neurons p = 0.0823. TTX, n = 52 radial astrocytes and 123 neurons from 7 animals, radial astrocytes ****p < 0.0001, neurons ****p < 0.0001.

### Radial astrocytes respond to visual stimulation with increases in calcium transients

We next tested whether the presentation of a behaviourally relevant visual stimulus^37^ could induce calcium transients in radial astrocytes in intact animals (Fig. 3). We imaged the cells of the optic tectum in darkness to establish a baseline of spontaneous activity in the circuit and then presented a looming stimulus every 6 s to the contralateral eye of the animal (Fig. 3a-d and Video 5). Compared to baseline, visual stimulation significantly increased both the number of active glia and neurons (Fig. 3e), while also significantly increasing the number of calcium events in both cell types (Fig. 3f). Repeated looming stimulus also significantly increased the mean number of events per cell for both cell types (Fig. 3g,h). Together this further demonstrates that the calcium activity of glia in the optic tectum is coupled to neuronal activity and shows that radial astrocytes in *Xenopus* respond to visual stimulation early in development.

**Fig. 3:**
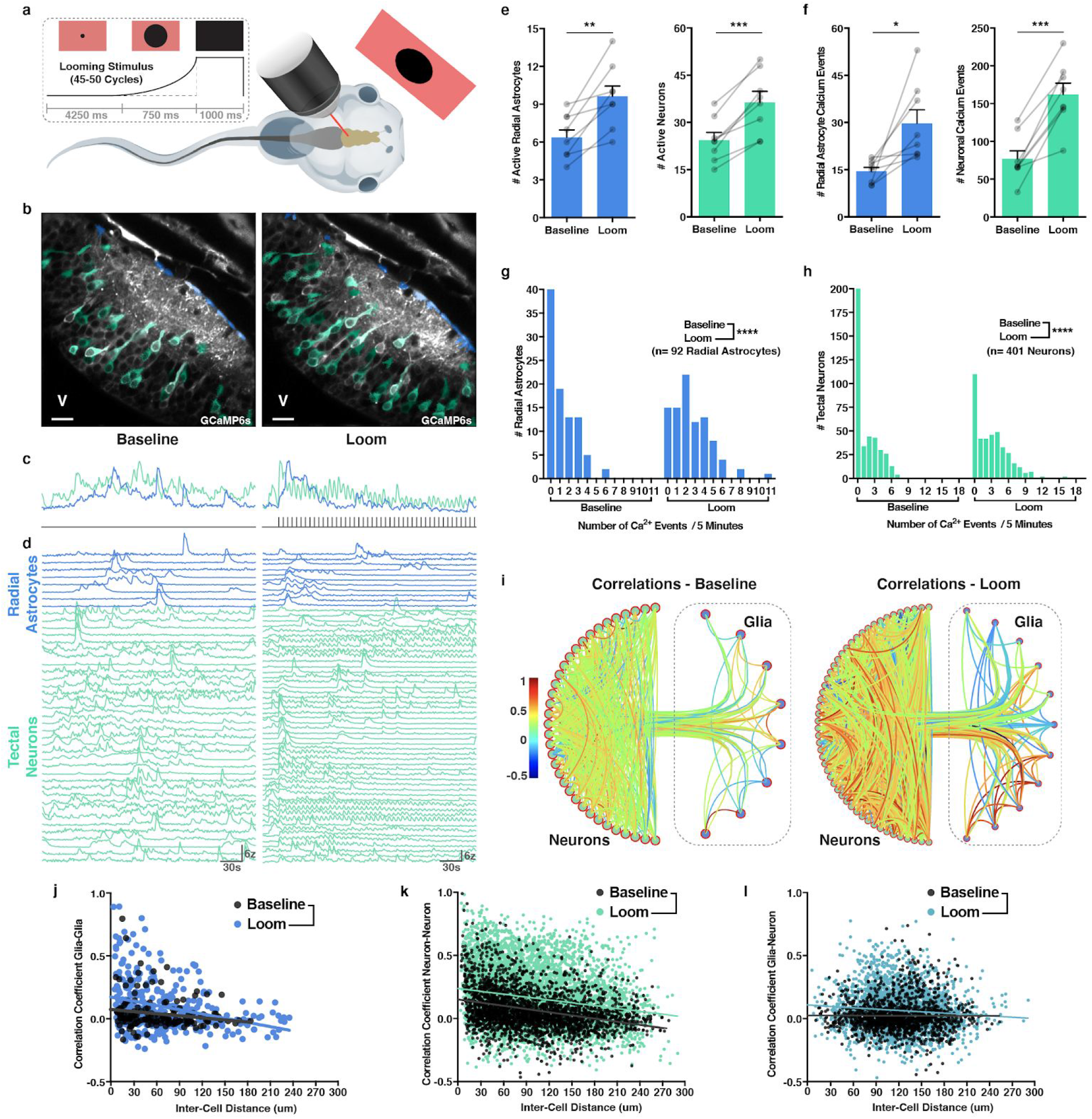
Repeated visual stimulation drives Ca^2+^ activity in radial astrocytes and induces correlation between neighbouring cells. **a**, Schematic of a *Xenopus laevis* tadpole being presented a repeated looming stimulus during live imaging of cells in the optic tectum. Looming stimulus presented every 6 s, after 4250 ms, a small dark dot expands over 750 ms and fills the screen for 1000 ms before resetting and repeating. **b**, Average temporal projections, 4500 frames (t), baseline (left), looming stimulus (right) (V= ventricle, scale bar = 25 μm). **c**, Average traces of Ca^2+^ activity during baseline (left) and looming stimulus (right), radial astrocytes (blue), neurons (green), looming stimulus presentation (grey lines), total time = 300 s. **d**, Traces of resting state and evoked Ca^2+^ activity during baseline (left) and looming stimulus (right), radial astrocytes (blue), neurons (green), total time = 300 s. **e**, Number of active cells during each imaging period, radial astrocytes (blue), neurons (green) (coloured bars = mean + s.e.m.). Paired t-tests, n = 8 animals, radial astrocytes **p = 0.0012, neurons ***p = 0.0002. **f**, Total number of Ca^2+^ events during each imaging period, radial astrocytes (blue), neurons (green) (coloured bars = mean + s.e.m.). Paired t-tests, n = 8 animals, radial astrocytes *p = 0.0145, neurons ***p = 0.0001. **g**, **h**, Number of events per cell during each imaging period, radial astrocytes (blue), neurons (green). Mann-Whitney tests, n = 92 glia and 401 neurons from 6 animals, (**h**) radial astrocytes ****p < 0.0001,(**i**) neurons ****p < 0.0001. **i**, Circular graphs showing correlations between neurons (left half), between radial astrocytes (right half), and between neurons and radial astrocytes (linkage between half-circles), baseline (left), looming stimulus (right). Cell order reflects relative locations of cells. **j**, Correlation vs. distance plot, radial astrocytes, baseline (grey dots), looming stimulus (blue dots), n = 481 pairs from 8 animals, difference between slopes p = 0.1453, difference between elevations **p = 0.0011. **k**, Correlation vs. distance plot, neurons, baseline (grey dots), looming stimulus (green dots), n = 8500 pairs from 8 animals, difference between slopes p = 0.3465, difference between elevations, ****p < 0.0001. **l**, Correlation vs. distance plot, neuron-radial astrocyte, baseline (grey dots), looming stimulus (turquoise dots), n = 4360 pairs from 8 animals, difference between slopes **p = 0.0025.

### Visually-evoked calcium responses in radial astrocytes exhibit delayed onsets and peak

To better understand the relationship between neuronal and glial responses to visual stimulation, we quantified the delay between the onset of neuronal events and those in glia, the delay between the peak of averaged neuronal and glial responses, and the number of looming stimulus events it took for both cell types to reach a peak response (Fig. 3c). On average there was a 4.35 ± 1.70 s delay between the onset of neuronal responses and those in glial cells, and a 15.43 ± 4.08 s delay between the peak of the neuronal responses and the peak of the responses in glia. Neurons reached a peak response after 1.43 ± 0.30 looming stimuli while the peak of the averaged glial responses occurred after 3.86 ± 0.80 presentations of the looming stimulus. This suggests that while glia do respond to individual visual events, at a circuit level, the collective response of glia to visual activity is tuned more towards integrating visual information over time.

### Visual stimulation enhances correlation between neighbouring radial astrocytes

We next assessed measures of temporal and spatial correlation between calcium responses in all cell types in order to gain information about functional connectivity in the optic tectum. Pearson correlation coefficients were calculated using Brainstorm software^38^ for each pairwise combination of calcium traces. In each animal, visual stimulation increased the mean pairwise correlation coefficients between cells, both for neurons and for glia (Fig. 3i). To determine whether the correlation between cells is proportional to the distance between them, we pooled all of the correlation data from 8 animals and then plotted the correlation coefficients against the distance between each corresponding pair of cells (Fig. 3j,k,l). To quantify these relationships, we calculated simple linear regressions of the pairwise correlations, comparing baseline and visual stimulation conditions. During baseline conditions, neuron-neuron and glia-glia, but not neuron-glia correlations, were related to the distance between the cells (as measured by significantly non-zero negative slopes). During visual stimulation, the relationship between correlated activity and distance remained for neuron-neuron and glia-glia pairs and a previously undetected relationship emerged for the neuron-glia pairs. Visual stimulation also significantly increased mean correlations across glia-glia (baseline: 0.047±0.011, loom: 0.097±0.011, Fig. 3j) and neuron-neuron (baseline: 0.069±0.003, loom: 0.165±0.003, Fig.3k) pairs. Lower correlations were observed for glia-neuron pairs (baseline: 0.024±0.003, loom: 0.066±0.003, Fig.3l). Together these data indicate that the activity of both neurons and glia are influenced by sensory inputs to the optic tectum; however, the comparative lack of spatial clustering of correlated glia-neuron pairs in the tectum, even during visual stimulation, suggests that their activity is independently regulated.

### Activation of AMPARs or mGluR1, but not mGluR5, induces calcium transients in radial astrocytes

Despite extensive research into glial glutamate receptors and their roles in neuron-glial signaling, the identity and function of glial glutamate receptors that govern biologically relevant calcium signaling *in vivo* remain controversial. Given that RGC inputs into the tectum are understood to be exclusively glutamatergic^39^, we next sought to determine whether glutamate receptor agonists could induce responses in radial astrocytes during early development (Fig. 4 and Videos 6-9). Previous research in our lab has demonstrated that NMDARs are not present on radial astrocytes during these developmental stages^34,40^, so we focused on AMPARs and mGluRs, specifically mGluR1 and mGluR5 given the weight of the literature suggesting a role for these type-1 mGluRs during early development^16^. Compared to control treatment, AMPA (100 μM) in animals pretreated with TTX (1 μM) did not significantly increase the number of active glia but did increase the number of active neurons, while the mGluR1/5 agonist DHPG (200μM) showed a non-significant increase in the number of active glia and a significant increase in the number of active neurons. In radial astrocytes, AMPA and DHPG also significantly increased the mean number of calcium events per animal and per cell (Fig. 4p,q,r,). However, while treatment with AMPA significantly increased the mean number of events per active neuron (Fig. 4s), no comparable increase was observed in neurons treated with DHPG. Unexpectedly^41^, the mGluR5 specific agonist CHPG (200 μM) showed no increases in any measure of cellular activity for either neurons or glia (Fig. 4n-s). Together these data demonstrate that activation of AMPARs or mGluR1, but not mGluR5, can induce calcium transients in both tectal neurons and radial astrocytes.

**Fig. 4:**
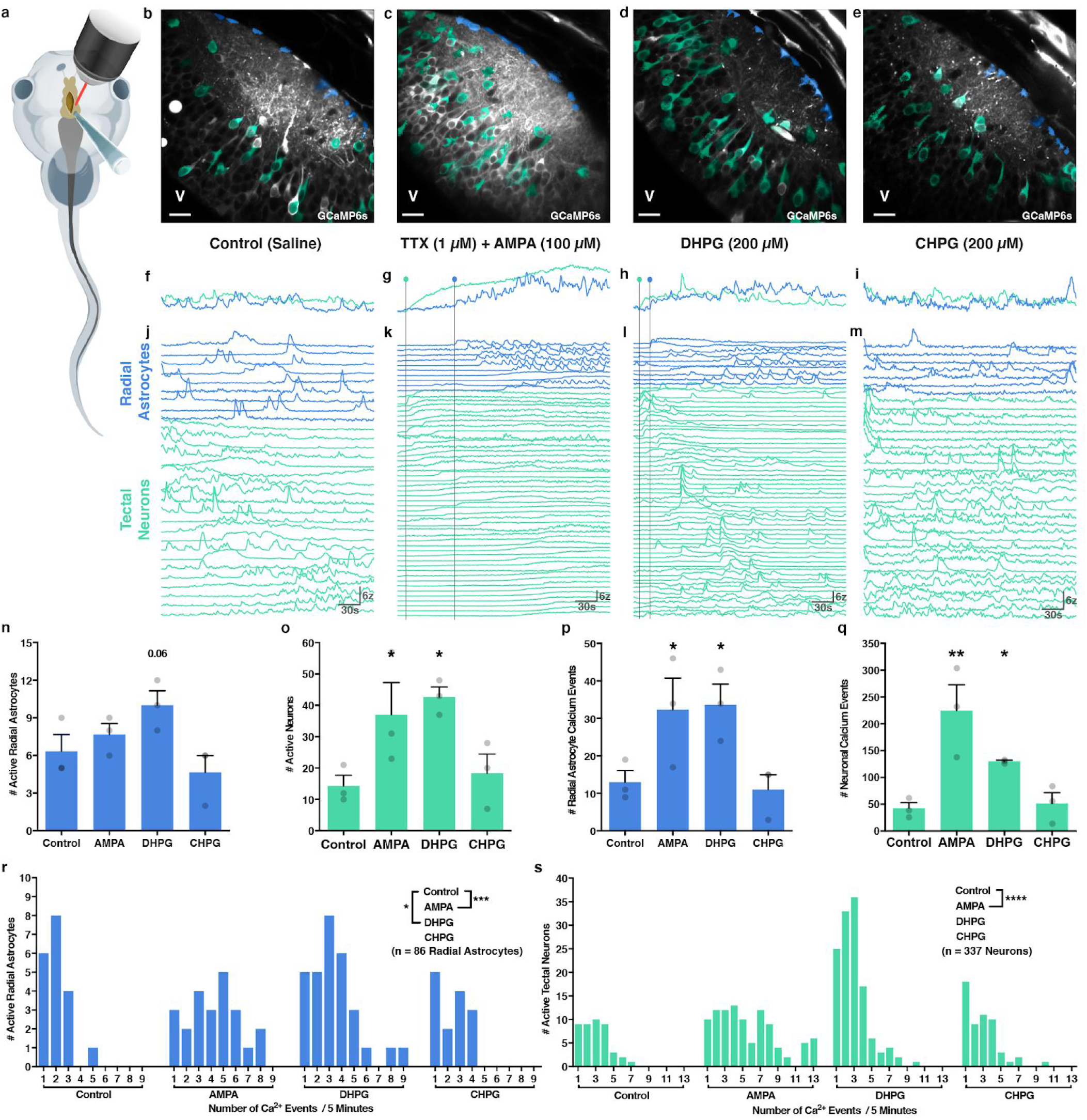
Activation of AMPARs or mGluR1 - but not mGluR5 - induces Ca^2+^ transients in tectal neurons and radial astrocytes during early development. **a**, Schematic of a *Xenopus laevis* tadpole with a slit cut into the ventricle to expose the optic tectum to allow acute pharmacological manipulation during live imaging. **b-e**, Average temporal projections, 4500 frames (t), external saline control (**b**), 1 μm TTX and 100 μM AMPA (**c**), 200 μM DHPG (**d**), 200 μM CHPG (**e**) (V= ventricle, scale bar = 25 μm). **f-i**, Average traces of resting state Ca^2+^ activity, external saline control (**f**), 1 μm TTX and 100 μM AMPA (**g**), 200 μM DHPG (**h**), 200 μM CHPG (**i**), radial astrocytes (blue), neurons (green), total time = 300 s. **j-m**, Traces of resting state Ca^2+^ activity, external saline control (**j**), 1 μm TTX and 100 μM AMPA (**k**), 200 μM DHPG (**l**), 200 μM CHPG (**m**), radial astrocytes (blue), neurons (green), vertical lines denote onset of drug action in neurons (green) and radial astrocytes (blue), total time = 300 s. **n**, **o**, Number of active cells during each imaging period, radial astrocytes (blue), neurons (green) (coloured bars = mean + s.e.m.). One-way ANOVAs, n = 3 animals per treatment group (12 total), (**n**) radial astrocytes F = 3.582 p = 0.0662, Fisher’s LSD tests, p AMPA = 0.4511, p DHPG = 0.0610, p CHPG = 0.3511, (**o**) neurons F = 4.676 *p = 0.0360, Fisher’s LSD tests, *p_AMPA_ = 0.0369, *p = 0.0141, p = 0.6707. **p**, **q**, Total number of Ca^2+^ events during each imaging period, radial astrocytes (blue), neurons (green) (coloured bars = mean + s.e.m.). One-way ANOVAs, n = 3 animals per treatment group (12 total), (**p**) radial astrocytes F = 4.689 *p = 0.0358, Fisher’s LSD tests, *p_AMPA_ = 0.0410, *p_DHPG_ = 0.0315, p_CHPG_ = 0.8076, (**q**) neurons F = 10.12 **p = 0.042, Fisher’s LSD tests, *p_AMPA_ = 0.0013, *p_DHPG_ = 0.0484, p_CHPG_ = 0.8172. **r**, **s**, Number of events per cell during each imaging period, radial astrocytes (blue), neurons (green). One-way ANOVAs, n = 86 radial astrocytes and 337 neurons from 12 animals (3 per treatment), (**r**) radial astrocytes F = 6.667 ***p = 0.0004, Dunnett’s multiple comparison test, ***p _AMPA_ = 0.0003, *p _DHPG_ = 0.0292, p _CHPG_ = 0.9212, (**s**) neurons F = 29.76 ****p < 0.0001, Dunnett’s multiple comparison test, ****p _AMPA_ < 0.0001, p _DHPG_ = 0.9969, p _CHPG_ = 0.9914.

### Delay in responses between radial astrocytes and tectal neurons to glutamate receptor agonists suggests indirect activation of glial cells

Further analysis into the temporal kinetics of the responses of tectal cells to AMPA and mGluR1 agonists revealed a stark contrast in the time between activation of tectal neurons and radial astrocytes. Accounting for the rate of diffusion of the drugs through the tectum, it was apparent that neuronal responses precede glial responses by a substantial margin. The first calcium responses in glia initiated by AMPA and DHPG occured 76.2 ± 7.9 s and 16.0 ± 4.2 s respectively following calcium rises in neurons occupying the same spatial regions of the optic tectum (Fig. 4k,l). We also demonstrated that the delay between tectal neuron and radial astrocyte activation during visual stimulation has a mean lag time of 4.35 ± 1.70 s, a delay much shorter than that of agonist induced responses (Fig. 3). Taken together, these results suggest that calcium increases in radial astrocytes in response to AMPAR and mGluR1 activation may be indirect, possibly occurring downstream of postsynaptic neuronal activation, and due to the mismatch in temporal kinetics of the responses, these glutamate receptors are unlikely to be the main mediators of visually induced calcium events in radial astrocytes.

### Broad spectrum blockade of glutamate receptors in the optic tectum enhances radial astrocyte responses to visual stimulation

Given that activation of AMPARs or mGluR1 both caused increases in calcium transients in radial astrocytes, we next sought to understand the role these glutamate receptors play in contributing to sensory-evoked calcium transients in glia (Fig. 5). To do this, we treated animals with a cocktail of pharmacological agents to block all glutamate receptors in the optic tectum (NMDARs - 100 μM R-CPP; AMPA/KainateRs - 50 μM DNQX, mGluRs - 100 μM LY341495^42^; mGluR1 - 100 μM CPCCOEt) and then presented a looming stimulus to the contralateral eye every 6 s (Fig. 5a). As expected, tectal neurons exhibited a dramatic reduction in spontaneously occurring calcium transients, and no longer responded to the looming stimulus (Fig. 5b-p, Fig. 3c,d,e,f,h). Surprisingly, during glutamate receptor blockade, radial astrocytes exhibited a significant increase in spontaneously occurring calcium transients and continued to significantly respond to the presentation of the looming stimulus (Fig. 5k,m,o and Video 10). These results imply the presence of a signaling pathway independent of ionotropic and metabotropic glutamate receptors capable of mediating visually-evoked calcium transients in radial astrocytes, and it suggests that radial astrocytes are directly responsive to presynaptic release from RGC axon terminals.

**Fig. 5:**
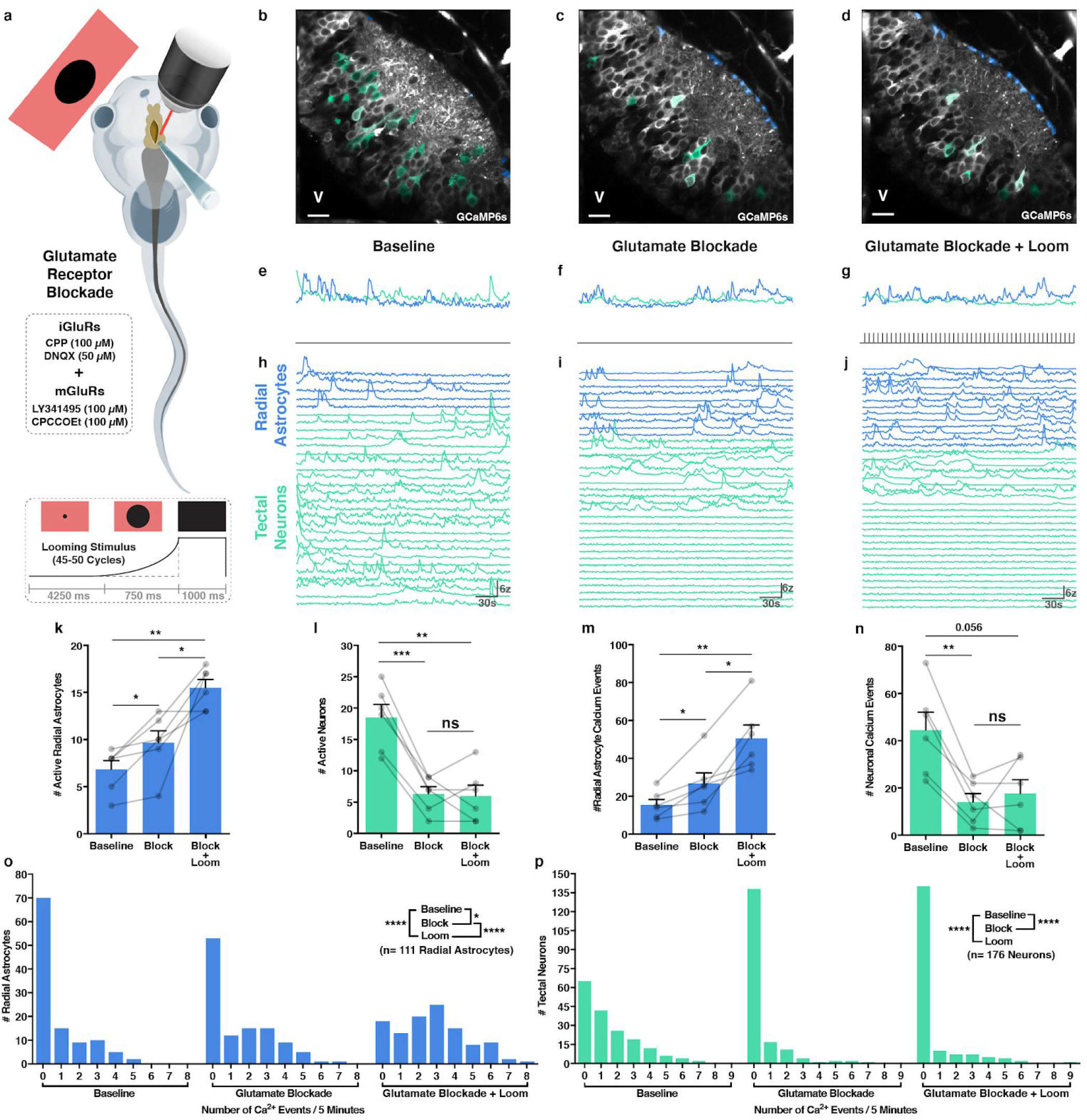
Blockade of ionotropic and metabotropic glutamate receptors in the optic tectum enhances both spontaneous and visually evoked Ca^2+^ activity in radial astrocytes. **a**, Schematic of a *Xenopus laevis* tadpole with a slit cut into its ventricle to expose the optic tectum to acute pharmacological manipulation being exposed to repeated looming stimulus presentation during live imaging. Glutamate receptors blockade cocktail consisted of 100 μM CPP, 50 μM DNQX, 100 μM LY341495, and 100 μM CPPCCOEt. Looming stimulus presented every 6 s. **b-d**, Average temporal projections, 4500 frames (t), baseline (**b**), after glutamate receptor blockade (**c**), looming stimulus presentation (**d**) (V= ventricle, scale bar = 25 μm). **e-g**, Average traces of resting state and evoked Ca^2+^ activity, baseline (**e**), after glutamate receptor blockade (**f**), looming stimulus presentation (**g**), radial astrocytes (blue), neurons (green), total time = 300 s. **h-j**, Traces of resting state and evoked Ca^2+^ activity, baseline (**h**), after glutamate receptor blockade (**i**), looming stimulus presentation (**j**), radial astrocytes (blue), neurons (green), looming stimulus presentation (grey lines), total time = 300 s. **k**, **l**, Number of active cells during each imaging period, radial astrocytes (blue), neurons (green) (coloured bars = mean + s.e.m.). One-way repeated measures ANOVAs, n = 6 animals, (**k**) radial astrocytes F = 21.56 **p = 0.0027, Tukey’s multiple comparisons test, baseline vs. block *p = 0.0428, baseline vs. loom **p = 0.0023, block vs. loom *p = 0.0486, (**l**) neurons F = 39.76 ****p < 0.0001, Tukey’s multiple comparisons test, baseline vs. block ***p = 0.0004, baseline vs. loom **p = 0.0027, block vs. loom p = 0.9778. **m**, **n**, Total number of Ca^2+^ events during each imaging period, radial astrocytes (blue), neurons (green) (coloured bars = mean + s.e.m.). One-way repeated measures ANOVAs, n = 6 animals, (**m**) radial astrocytes F = 26.09 ***p = 0.0005, Tukey’s multiple comparisons test, baseline vs. block *p = 0.0465, baseline vs. loom **p = 0.0039, block vs. loom *p = 0.0164, (**n**) neurons F = 12.58 **p = 0.0070, Tukey’s multiple comparisons test, baseline vs. block **p = 0.0043, baseline vs. loom p = 0.0565, block vs. loom p = 0.8079. **o**, **p**, Number of events per cell during each imaging period, radial astrocytes (blue), neurons (green). One-way repeated measures ANOVAs, n = 111 radial astrocytes and 176 neurons from 6 animals, (**o**) radial astrocytes F = 37.00 ****p < 0.0001, Tukey’s multiple comparisons test, baseline vs. block *p = 0.0190, baseline vs. loom ****p < 0.0001, block vs. loom ****p < 0.0001, (**p**) neurons F = 26.83 ****p < 0.0001, Tukey’s multiple comparisons test, baseline vs. block ****p < 0.0001, baseline vs. loom ****p < 0.0001, block vs. loom p = 0.6992.

### Blockade of EAATs inhibits visual stimulation-induced calcium transients in radial astrocytes

A number of experiments have suggested a role for EAATs in contributing to calcium transients in glia^15,18^. Because we found that glutamate receptor blockade was able to isolate the calcium response of glia from any significant postsynaptic neuronal activity, we again blocked all glutamate receptors in the optic tectum while also testing the effects of blocking the principal routes of glutamate reuptake by glia through EAAT 1 and 2 by presenting visual stimulation in the presence of TFB-TBOA (500 nM) (Fig. 6a,b,c and Video 11). In contrast to glutamate receptor blockade alone, the additional blockade of EAATs resulted in a decrease in glial activation during visual stimulation, suggesting that glutamate reuptake mediates calcium transients in radial astrocytes in response to vision, independent of postsynaptic neuronal activation (Fig. 6d,e,f,g). However, glutamate reuptake by EAATs is not thought to directly import calcium into the cell, indicating that the calcium response is likely mediated by downstream signaling partners.

**Fig. 6:**
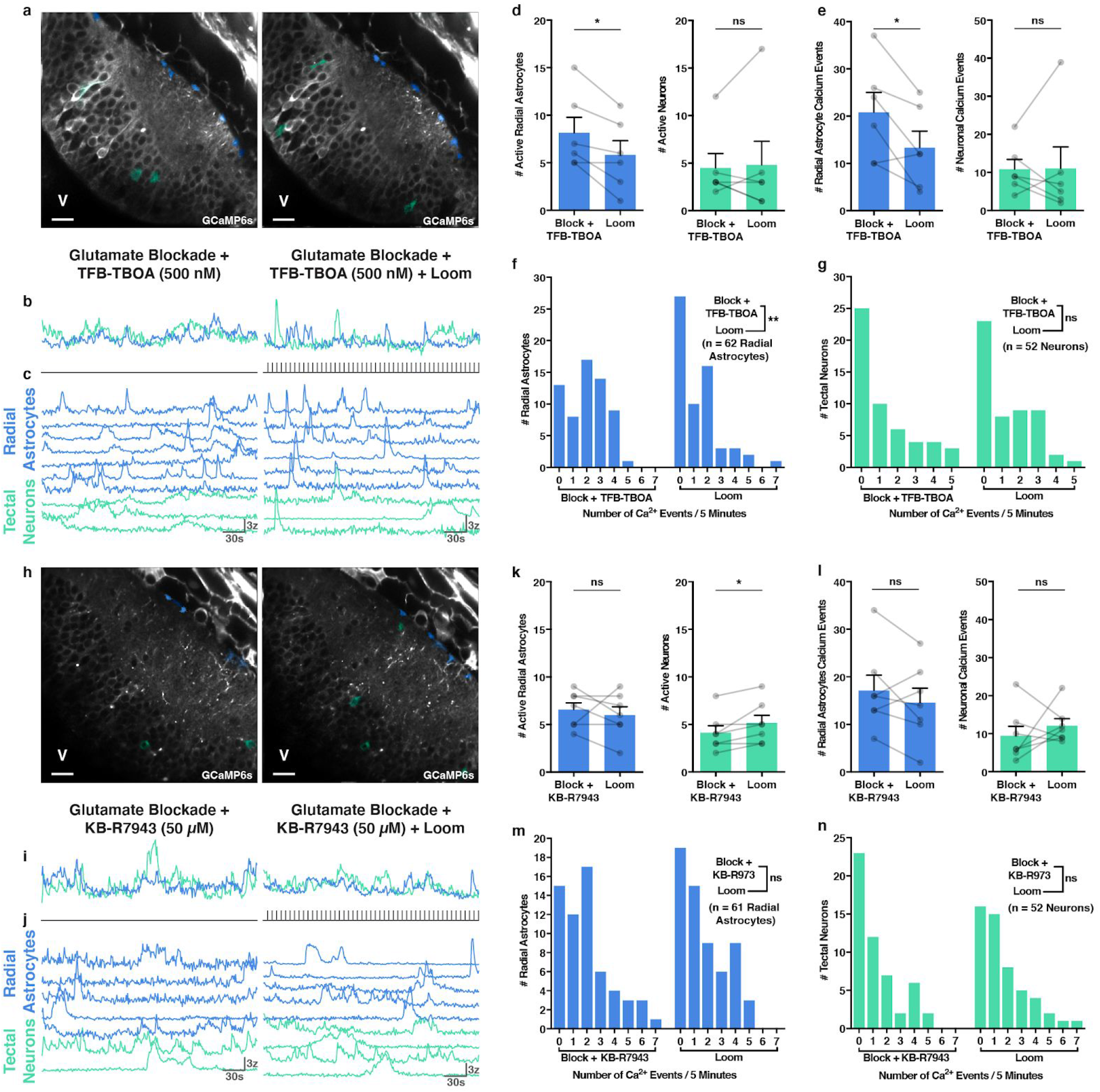
Visually evoked Ca^2+^ transients in radial astrocytes require glutamate reuptake through EAATs and reversal of the sodium-calcium exchanger. **a**, Average temporal projections, 4500 frames (t), glutamate receptor blockade and 500 nM TFB-TBOA (left), looming stimulus presentation (right) (V= ventricle, scale bar = 25 μm). **b**, Average traces of resting state and evoked Ca^2+^ activity, baseline (**e**), after glutamate receptor blockade and 500 nM TFB-TBOA (left), looming stimulus presentation (right), radial astrocytes (blue), neurons (green), total time = 300 s. **c**, Traces of resting state and evoked Ca^2+^ activity, after glutamate receptor blockade and 500 nM TFB-TBOA (left), looming stimulus presentation (right), radial astrocytes (blue), neurons (green), looming stimulus presentation (grey lines), total time = 300 s. **d**, Number of active cells during each imaging period, radial astrocytes (blue), neurons (green) (coloured bars = mean + s.e.m.). Paired t-tests, n = 6 animals, radial astrocytes *p = 0.0173, neurons p = 0.7771. **e**, Total number of Ca^2+^ events during each imaging period, radial astrocytes (blue), neurons (green) (coloured bars = mean + s.e.m.). Paired t-tests, n = 6 animals, radial astrocytes *p = 0.0312, neurons p = 0.9681. **f**, **g**, Number of events per cell during each imaging period, radial astrocytes (blue), neurons (green). Mann-Whitney tests, n = 62 radial astrocytes and 52 neurons from 6 animals, (**e**) radial astrocytes **p = 0.0014, (**f**) neurons p = 0.7403. **h**, Average temporal projections, 4500 frames (t), after glutamate receptor blockade and 50 μM KB-R7943 (left), looming stimulus presentation (right) (V= ventricle, scale bar = 25 μm). **I**, Average traces of resting state and evoked Ca^2+^ activity, baseline (**e**), after glutamate receptor blockade and 50 μM KB-R7943 (left), looming stimulus presentation (right), radial astrocytes (blue), neurons (green), total time = 300 s. **j**, Traces of resting state and evoked Ca^2+^ activity, after glutamate receptor blockade and 50 μM KB-R7943 (left), looming stimulus presentation (right), radial astrocytes (blue), neurons (green), looming stimulus presentation (grey lines), total time = 300 s. **k**, Number of active cells during each imaging period, radial astrocytes (blue), neurons (green) (coloured bars = mean + s.e.m.). Paired t-tests, n = 7 animals, radial astrocytes p = 0.5352, neurons *p = 0.0382. **l**, Total number of Ca^2+^ events during each imaging period, radial astrocytes (blue), neurons (green) (coloured bars = mean + s.e.m.). Paired t-tests, n = 7 animals, radial astrocytes p = 0.2626, neurons p = 0.4375. **m**, **n**, Number of events per cell during each imaging period, radial astrocytes (blue), neurons (green). Mann-Whitney tests, n = 61 radial astrocytes and 52 neurons from 7 animals, (**k**) radial astrocytes **p = 0.0014, (**l**) neurons p = 0.2152.

### Blockade of the sodium-calcium exchanger prevents visual stimulation-induced calcium transients in radial astrocytes

EAATs co-transport 3 sodium ions into the cell along with each molecule of glutamate. A number of studies have demonstrated that, at least *in vitro* or in *ex vivo* preparations, the sodium-calcium exchanger (NCX) on glia, which under normal circumstances uses electrochemical gradients to export calcium and import sodium, reverses direction of ion flow following rapid increases in internal sodium concentration resulting in the extrusion of sodium ions and import of extracellular calcium ions^32^. To determine whether this mechanism may also underlie visually-evoked glial calcium activity *in vivo*, we again blocked all glutamate receptors in the optic tectum while additionally blocking the reverse mode of NCX with 50 μM KB-R7943 during visual stimulation (Fig. 6h,i,j and Video 12). Blockade of the reverse mode of NCX produced results that mimicked the blockade of EAATs. No increase in glial calcium activity in response to visual stimulation could be elicited in the presence of the NCX blocker, suggesting that the activity of EAATs and NCX family members are coupled, likely via internal sodium concentration, thereby mediating calcium responses in glia during visual stimulation (Fig. 6k,l,m,n).

### Blockade of NCX alone is sufficient to prevent radial astrocytes from responding to visual stimulation

Given that blockade of NCX prevented radial astrocytes from responding to visual stimulation in the presence of glutamate receptor blockers that significantly reduced postsynaptic activity, we set out to determine whether impairing NCX function alone is sufficient to prevent radial astrocytes from responding to visual stimulation. We treated animals with 50 μM KB-R7943 alone before repeating our visual stimulation protocol and quantifying responses (Fig. 7a-f and Video 13). Blockade of reverse mode NCX did not prevent tectal neurons from responding to the looming stimulus although it did often alter their response properties in such a way as to produce progressively greater responses across time - consistent with reduced glutamate clearance (neuronal responses peaked after 17.5 ± 5.8 looms vs. 1.43 ± 0.3 looms without pharmacological intervention) (Fig. 7d,f,g-j). In contrast, radial astrocytes did not exhibit any increases in calcium activity during visual stimulation (Fig. 7g,h,i). This suggests that during development the visually-evoked calcium activity in radial astrocytes occurring *in vivo* is more dependent upon calcium influx through NCX following reuptake of glutamate than either direct or indirect activation of ionotropic or metabotropic glutamate receptors.

**Fig. 7:**
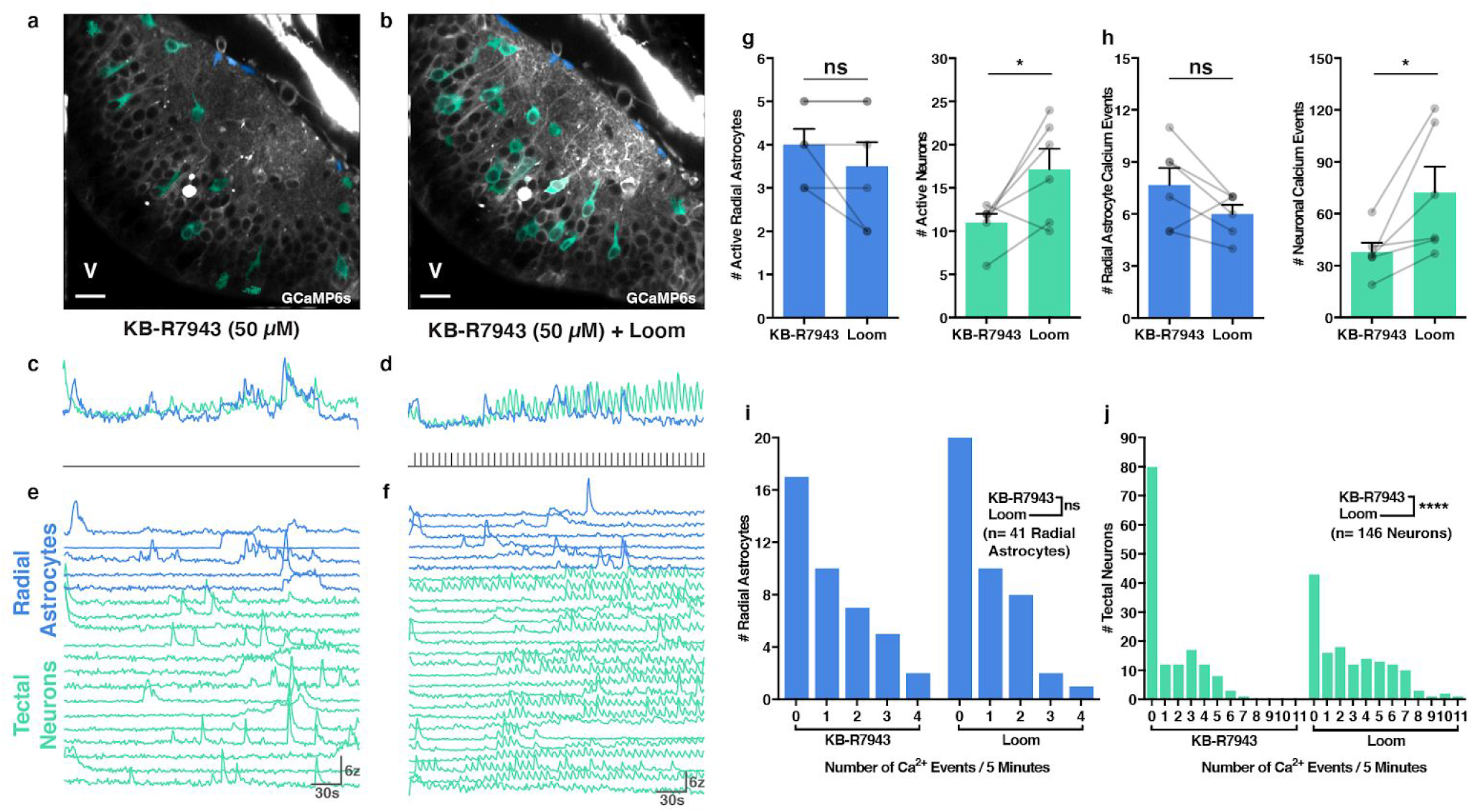
Blocking the reverse mode of NCX is sufficient to block the radial astrocyte response to visual stimulation. **a**, **b**, Average temporal projections, 4500 frames (t), 50 μM KB-R7943 (**a**), looming stimulus presentation (**b**) (V= ventricle, scale bar = 25 μm). **c**, **d**, Average traces of resting state and evoked Ca^2+^ activity, 50 μM KB-R7943 (**c**), looming stimulus presentation (**d**), radial astrocytes (blue), neurons (green), looming stimulus presentation (grey lines), total time = 300 s. **e**, **f**, Traces of resting state and evoked Ca^2+^ activity, 50 μM KB-R7943 (**e**), looming stimulus presentation (**f**), radial astrocytes (blue), neurons (green), total time = 300 s. **g**, Number of active cells during each imaging period, radial astrocytes (blue), neurons (green) (coloured bars = mean + s.e.m.). Paired t-tests, n = 6 animals, radial astrocytes p = 0.2031, neurons *p = 0.0421. **h**, Total number of Ca^2+^ events during each imaging period, radial astrocytes (blue), neurons (green) (coloured bars = mean + s.e.m.). Paired t-tests, n = 6 animals, radial astrocytes p = 0.1051, neurons *p = 0.0349. **i**, **j**, Number of events per cell during each imaging period, radial astrocytes (blue), neurons (green). Mann-Whitney tests, n = 41 radial astrocytes and 146 neurons from 6 animals, (**i**) radial astrocytes p = 0.3698, (**j**) neurons ****p < 0.0001.

## Discussion

While early research into glial calcium signaling suggested mGluRs directly mediate glial calcium events in response to physiological stimulation^14^, research over the last decade has begun to demonstrate that neurotransmitter transporters such as EAATs and Gamma-Aminobutyric acid transporters (GATs) mediate a diversity of glial calcium responses^15,27,28,31,43^. Developing a thorough mechanistic understanding of how glial cells detect and integrate sensory-evoked neurotransmission requires the implementation of methods that can assess the various contributions of, and interplay between, both receptor and transporter mediated signaling pathways simultaneously in both glia and neurons during sensory processing in awake animals. Through the combined use of visual stimulation to drive, and pharmacology to modulate, neural activity in the intact *Xenopus laevis* visual system, we have demonstrated that during early development radial astrocytes in the neuropil of the retinotectal circuit respond to visual stimulation through temporally-correlated increases in calcium activity which result from glutamate uptake through EAATs and subsequent calcium entry through NCX. Unexpectedly, sensory-evoked activity in these glia occurred independently of the activation of both ionotropic and metabotropic glutamate receptors. This expands on the accumulating literature showing that the activity of sodium-dependent neurotransmitter transporters and NCX are functionally connected^26,32^ by demonstrating a role for NCX in regulating sensory-evoked glial activity *in vivo* during early development.

In the visual cortex of adult mammals mature astrocytes have highly sensitive receptive fields that reflect the topographic organization of the visual system and respond to complex spatial components of visual activity such as orientation selectivity through a mechanism involving glutamate transporters^15^. Here we demonstrate that even during early development of the amphibian visual system, radial astrocytes are capable of responding to individual visual events. We observed that glia exhibit temporally correlated responses to visual stimulation which peak during periods of persistent visual activity, and that the temporal correlation of glial activity is highest between neighbouring glial cells. Together this suggests that in the developing optic tectum, glia are already responsive to visual information in a manner that reflects a primitive representation of the spatial organization of the visual system (Fig. 8a). Given that the mechanism underlying the calcium responses of these glia requires glutamate reuptake, and that responses peak during periods of persistent visual activity, it suggests that these cells are likely processing information related to how much presynaptic glutamate release is occurring across time and thus may also encode information relating to spatial components of the visual information. Experiments in adult mammals have demonstrated a similar role for glutamate transporters in visually-evoked calcium responses in adult astrocytes, demonstrating the importance of this mechanism throughout life. Glia may also be able to process higher order visual information through the integration of additional signaling mechanisms which may act independently of or modulate the pathway we have identified here. Additionally, in response to being activated by neuronal activity, glia are known to both modulate and negatively regulate neural activity through various mechanisms involving neurotransmitter transporters, neuromodulators, and release of gliotransmitters^10,12,21,28,31,44^. In our visual stimulation experiments we often observed a rundown in the calcium responses of tectal neurons to repeated visual stimulation (Fig.3c,d), and while here we cannot rule out non-glial mediated effects of pharmacological blockade of NCX in the optic tectum, when blocking the function of NCX, and as such reducing glial activity, we frequently observed neuronal responses which progressively increased in intensity throughout the period of visual stimulation (Fig.7d,f). While this effect could be mediated through multiple mechanisms, it is consistent with NCX blockade impairing glutamate reuptake through EAATs leading to the progressive accumulation of glutamate in the neuropil, or radial astrocytes playing an active role in negatively regulating neural activity as has been shown in zebrafish^12^. It will be valuable to mechanistically dissect glia to neuron signals occurring in the retinotectal circuit and determine how they modulate visual processing.

**Fig. 8:**
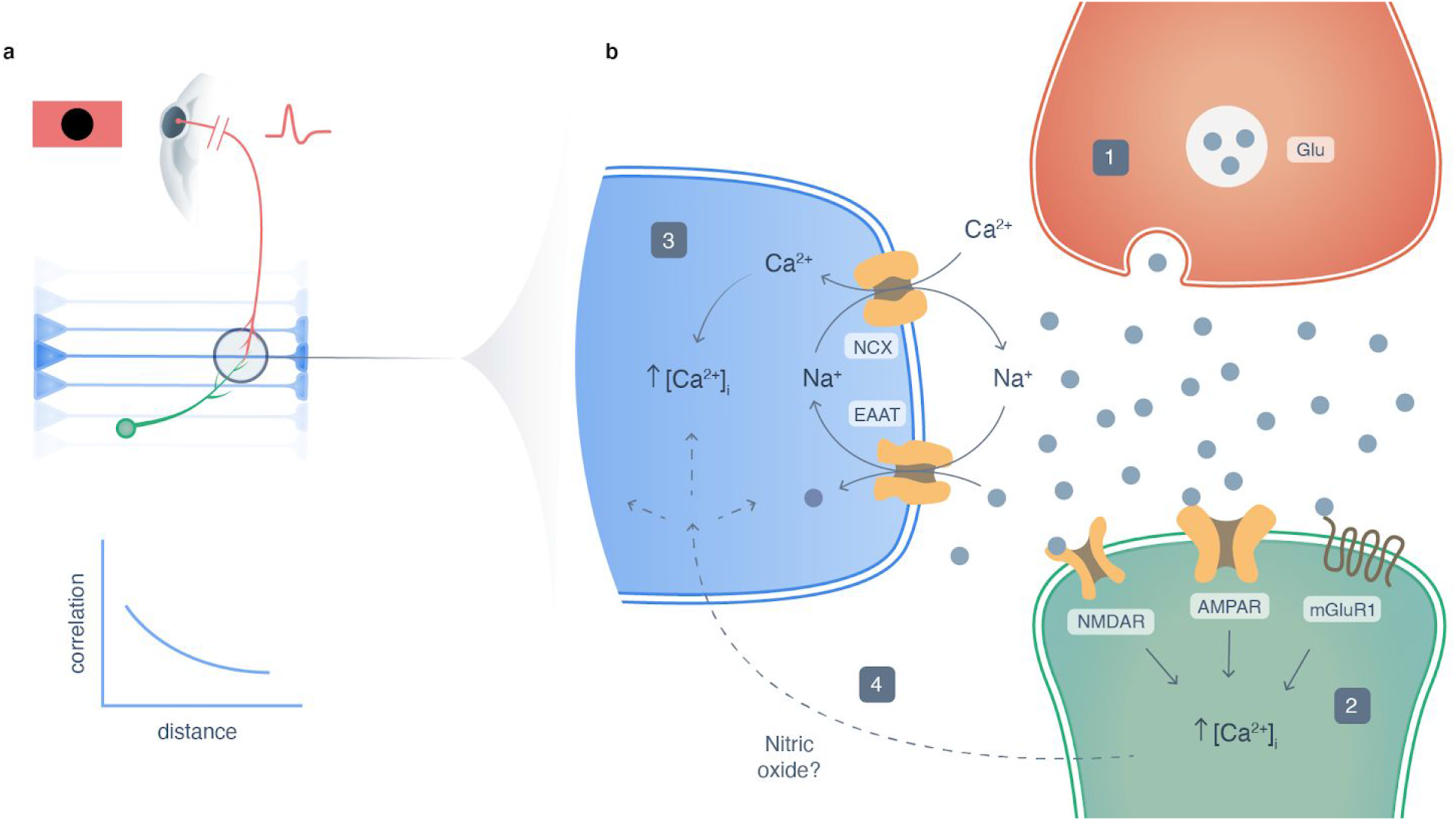
Sodium-calcium exchanger mediates sensory-evoked radial astrocyte calcium transients in the developing retinotectal system. **a**, Schematic showing a looming stimulus inducing temporally correlated activity in cells occupying the same region of the neuropil in the *Xenopus laevis* retinotectal circuit (retinal ganglion cell - red, radial astrocytes - blue, tectal neuron - green). **b**, Schematic of the mechanism underlying visually-evoked calcium activity in radial astrocytes of the retinotectal circuit. (1) Presynaptic release of glutamate from RGC axon terminals during visual stimulation induces (2) calcium elevations in tectal neurons through the activation of postsynaptic glutamate receptors followed by (3) calcium elevations in radial astrocytes through reversal of NCX and calcium influx following uptake of glutamate through EAATs. Activation of tectal neuron glutamate receptors and (4) the release of diffusible signals such as nitric oxide may modulate radial astrocyte calcium activity over larger timescales. Credit: Artwork, A. Desaulniers, Orcéine.

The temporal kinetics of the calcium responses occurring during visual stimulation in these radial astrocytes match closely with those observed in astrocytes in the visual cortex of adult ferrets^15^ and more recently in the somatosensory cortex of adult mice^44^, where sensory-evoked calcium responses lag behind neuronal responses by seconds. This suggests the possibility that a shared mechanism may be underlying these responses. Despite EAATs and NCX mediating visually driven calcium events in radial astrocytes, similarly to astrocytes in mammalian systems, these glia do also exhibit calcium rises in response to pharmacological activation of AMPARs and mGluR1. However, these agonist-induced events occur over timescales outside the physiological timescales over which calcium events occur in response to visual stimulation (AMPA: 76.2 ± 7.9 s, DHPG: 16.0 ± 4.2 s, visual stimulation: 4.35 ± 1.70 s). This suggests that the calcium events occurring in glia in response to activation of these receptors are not likely resulting from direct activation of these receptors on glia but rather may occur in response to the release of diffusible signals following postsynaptic activation of tectal neurons, as glutamate receptor activation is coupled to the release of diffusible signals such as nitric oxide and endocannabinoids. Despite glutamate receptors not mediating visually-driven calcium events in radial astrocytes, it remains possible the release of diffusible signals following activation of these receptors on neurons may still contribute to the regulation of EAAT/NCX-mediated calcium signaling in glia.

Glial calcium responses in our experiments often desensitized to the visual stimulus within the first couple of minutes of repeated stimulation, and consistent with the mechanism we have proposed above, it has been observed elsewhere that the activity-dependent internalization of the glial glutamate transporter GLT-1 requires calcium entry through NCX^45^. This suggests that visual stimulation may lead to the internalization of EAATs here as well. Nitric oxide and cGMP-dependent protein kinase (PKG) signaling, which we have previously demonstrated contribute to calcium signaling in radial astrocyte cell bodies and motility of their filopodial processes^34,46^, have also been shown to promote insertion of EAATs into the plasma membrane of Bergmann glia through a signaling cascade involving NO, cGMP, PKG, and calcium entry through NCX^47^. As such, the potential exists for postsynaptic activation of tectal neuron glutamate receptors and release of nitric oxide to facilitate insertion of EAATs into the glial membrane thereby preventing desensitization of glial calcium signaling in areas of the neuropil experiencing high levels of synaptic activity. Together this suggests that calcium entry through glial NCX may regulate glutamate clearance from active synapses. Whether prolonged alterations in glial NCX activity is capable of influencing neural circuit development remains an interesting open question.

Altogether our experiments demonstrate that during early development radial astrocytes are highly responsive to sensory stimulation *in vivo*. Mechanistically, visually-evoked calcium activity in these cells is driven by presynaptic release of glutamate, activation of EAATs and consequently calcium influx through NCX (Fig. 8b). Our experiments demonstrate that blockade of NCX alone is sufficient to prevent visually-evoked calcium activity in glia, and that the calcium events in these glia driven by sensory stimulation were not dependent to any significant degree on the activation of glutamate receptors. We have proposed a model that takes into account both the observations in the literature that glutamate receptors can mediate glial calcium signaling and our observations of glutamate receptor-independent calcium events. In conclusion, we have increased our understanding of a mechanism underlying glial responses to sensory stimulation *in vivo* by showing that, at least during early development, NCX mediates the calcium signal associated with glutamate reuptake through EAATs and that the temporal kinetics of these responses are equivalent to those observed in mature astrocytes in both mammalian visual^15^ and somatosensory cortex^44^. Our study further suggests that NCX may mediate important contributions of glia to neural circuit development, function, plasticity, and behaviour, relevant for both healthy function and disease.

## Methods

### Animal husbandry statement

Adult albino *Xenopus laevis* frogs were housed in the Montreal Neurological Institute animal care facility Centre for Neurological Disease Models. All animal use and experiments were approved by the Montreal Neurological Institute Animal Care Committee and in accordance with Canadian Council of Animal Care guidelines.

### GCaMP6s mRNA synthesis

The coding sequence of each GCaMP6s and mCherry were cloned into pCS2+ and the plasmids were linearized with NotI. Capped mRNA of GCaMP6s and mCherry were transcribed with the SP6 mMessage mMachine Kit (Ambion, Thermo Fisher). The GCaMP6s and mCherry mRNA were injected together at the two-cell stage into one blastomere of the embryo.

### Generation of GCaMP6s-expressing tadpoles

*In vitro* fertilization was used to prepare eggs for GCaMP6s mRNA injection. Adult female frogs were primed for egg laying by injection of pregnant mare serum gonadotropin (PMSG, 50UI, Prospec Bio) 4 days before *in vitro* fertilization. In the afternoon preceding *in vitro* fertilization, the frogs previously injected with PMSG were injected with human chorionic gonadotropin (hCG, 400UI, Sigma) to induce egg laying. The following morning an adult male frog was anesthetized by submersion in tricaine mesylate (MS-222, 0.2% w/v, Sigma) until reflexes were unresponsive (approximately 30 minutes). Once anesthetized, the head of the frog was severed and the skin on the belly cut open to expose internal organs. Testis were located and surgically removed and placed in ice chilled 1X Modified Barth’s Solution with HEPES (MBSH). Adult female frogs injected with hCG the previous day were then squeezed and eggs were collected in a clean 100 mm petri dish. A testis was placed in a 50 μL drop of 1X MBSH in the petri dish with the unfertilized eggs and then a portion of it was removed and macerated with a razorblade. A pipette tip was then used to mix the eggs with the macerated testis solution. To activate the sperm and initiate *in vitro* fertilization, 750 μL of 0.1X MBSH was added to the eggs and the mixture was kept at room temperature for 5 min before being submerged completely with 0.1X MBSH. After 10 min, embryos were degellied in cysteine (2% w/v in distilled water, pH 8.0, Sigma), washed twice in distilled water, then washed twice in 0.1X MBSH, and then monitored until they completed their first cell division and reached the two-cell stage. At this point, the embryos were transferred into a petri dish lined with a custom-made grid filled with 2% ficoll in 1X MBSH. Then 500 pg GCaMP6s mRNA along with 250 pg mCherry mRNA, in 2 nL RNAse-free water were pressure injected into one of the two cells of the embryo using a calibrated glass micropipette attached to a PLI-100 picoinjector (Harvard Apparatus). After injection the embryos in 2% ficoll in 1X MBSH were placed in an incubator and kept at 18 degree Celsius for 3 h before being transferred to 1% ficoll in 0.1X MBSH and then kept at 18 degrees Celsius overnight. The next morning embryos were transferred to 0.1X MBSH and raised at 18 degrees Celsius in a biological oxygen demand incubator with a 12 h light / 12 h dark cycle until they reached stage 47.

### Tectal electroporation of individual radial astrocytes

To visualize the structure of individual radial astrocytes, the optic tectum of stage 45 tadpoles was electroporated with plasmids encoding EGFP-F following protocols described previously by our lab. Tadpoles were anesthetised in MS-222 (0.02% in 0.1X MBSH), then a brief interventricular injection was performed using a glass micropipette containing 0.5-1 μg/uL pEGFP-F plasmid and the visual indicator fast green. After placing two platinum electrodes on either side of the optic tectum, a Grass Instruments SD9 electrical stimulator outfitted with a 3 μF capacitor was set to 37 V and used to apply three 1.6 ms pulses across the brain in each direction of polarity. Animals recovered in 0.1X MBSH for 48 hours before imaging.

### Preparing GCaMP6s-expressing tadpoles for live imaging

Stage 46-47 GCaMP6s-expressing tadpoles were screened for adequate levels of GCaMP6s expression on a Olympus BX-43 epifluorescence microscope prior to imaging. Animals were then immobilized by immersion in a solution of 2 mM pancurionium dibromide (Tocris) in 0.1X MBSH for 2-5 min. Then animals were placed in the middle of a 6 cm diameter petri dish lid and embedded in 0.8% low melting point agarose (Thermo Fisher). To bypass the blood brain barrier and permit for acute pharmacological intervention, a small incision was made into the dorsal surface of the skin of some tadpoles in between the hemispheres of the optic tectum using a 30 ga syringe needle to expose the associated ventricle. After embedding, intact animals were submerged in 9 mL of 0.1X MBSH, while those that underwent incision, were submerged in 9 mL of external saline solution (in mM – 115 NaCl, 2 KCl, 3 CaCl2, 3 MgCl2, 5 HEPES, and 10 glucose, pH 7.20, 250 mOsm).

### Two-photon microscope

Live imaging was performed using a Thorlabs multiphoton resonant scanner microscope with a 20X water-immersion objective (1.0 NA) mounted on a piezoelectric focus mount (PI). Fluorescence excitation light was generated using a Spectra-Physics InSight3x infrared femtosecond pulsed laser. Data was collected using ThorImage LS software.

### In vivo calcium imaging of GCaMP6s-expressing tadpoles

Before imaging, animals were left to settle under the microscope for 15 min to reduce drift in x, y, and z dimensions. Calcium imaging was then carried out using an excitation wavelength of 910 nm at an approximate power of 125 mW, measured before the scanhead. Each imaging epoch consists of 4500 images collected at 15 frames per second from a single z plane with x-y dimensions of 224.256 μm by 224.256 μm at a resolution of 512 by 512 pixels. Using ImageJ, images were then saved as tiff stacks for further processing.

Baseline timepoints were collected in darkness. For timepoints involving pharmacological treatments, compounds were first dissolved to the appropriate concentration in 1 mL of external saline solution and introduced to the bath in which the animal was embedded via micropipette. Agonists were applied to the bath immediately before an imaging epoch commenced while antagonists were applied immediately following the acquisition of baseline timepoints and allowed to equilibrate for 15 min before imaging. Visual stimuli were presented to the contralateral eye of the animal during treatment timepoints using a flat panel display screen (800 x 480, Adafruit, NY) covered with a red-filter (Wratten #29, Kodak) to prevent activation of the green channel PMT.

### Visual stimulation paradigm

For visual stimulation of GCaMP6s-expressing tadpoles, a video of a dark looming stimulus was created using Adobe After Effects (Video 14). A small dark dot remains stationary in the middle of the screen for 4250 ms before rapidly expanding to fill the visual field over 750 ms; the screen then remains dark for 1000 ms before resetting and repeating 45-50 times during the 5-minute treatment period.

### Registration and ROI extraction of raw calcium imaging data using Suite2p

Suite2p software (https://github.com/MouseLand/suite2p) was run using Python 3. Tiff stacks of raw calcium imaging data collected on a two-photon microscope were imported into Suite2p for registration and automated detection of regions of interest (ROIs) based on regions of dynamic calcium activity. Both rigid and nonrigid registration were performed using the default configurations of the software. Automated ROI detection used the default parameters of the software except the threshold scaling value was increased to 3.5 to favour detection of structures with a high signal to noise ratio. Then using the software GUI, ROIs were curated by hand to remove those that were neither tectal neuron cell bodies nor radial astrocyte endfeet. Raw fluorescence traces for each ROI were extracted from Suite2p as CSV files and used for further analysis.

### Baseline normalization of calcium traces and quantification of calcium events

To estimate the baseline fluorescence intensity of the calcium trace for each ROI, we calculated the bottom 5th percentile of values occurring in the calcium trace for that ROI during each 5-minute epoch. To quantify the number of calcium events occurring in an ROI during each 5 min epoch, we z-scored the calcium traces by subtracting the baseline from each value and then dividing by the standard deviation of the values in that trace. Increases in a calcium trace were only counted as events if they exceeded a z-score of 3.

### Correlation analysis, quantification, and visualization using Brainstorm

Suite2p files containing the ROIs and fluorescence traces for each experiment were imported into Brainstorm software^38^. Brainstorm was used to assess aspects of temporal correlation and functional connectivity between cells in the network while at rest and during visual stimulation. First, after importing the data, ROIs were annotated by hand as either being tectal neurons or radial astrocytes and segmented into left and right hemispheres respectively for later visualization. Then an N x N correlation matrix of Pearson correlation coefficients was calculated from all the pairwise combinations of ROIs from each animal. For each animal, network connectivity during each imaging epoch was then visualized by plotting the raw correlation matrix as a circular graph with tectal neuron ROIs along the left side, radial astrocyte ROIs along the right side, and the strength of the correlations between cells as colour-coded lines between ROIs.

To quantify the average level of correlation (both positive and negative) that exists between cells of each type in the network for each animal and whether it changes during visual stimulation, raw correlation matrices containing Pearson correlation coefficients for each pair of ROIs were first converted to absolute values. These matrices of absolute values were then analyzed by looking at each type of possible intercellular correlation (neuron - neuron, glial - glial, and neuron - glial). For each animal the mean Pearson correlation coefficient (absolute value) was calculated between all of its tectal neurons, between all of its radial astrocytes, and between all of its tectal neurons and radial astrocytes. Average correlation coefficients for each animal were then compared between resting state and visual stimulation.

To quantify whether the correlations that exist between cell types in the optic tectum are proportional to the distance between the cells and whether this relationship is influenced by visual stimulation, first we pooled the data from 8 animals into two groups: baseline and visual stimulation. We then extracted the linear distances between each of these pairs of ROIs using a custom Matlab script and matched them to the corresponding Pearson correlation coefficient for that pair. We processed the data in groups based on cell type like above by plotting intercellular distance against raw Pearson correlation coefficient for each pair of ROIs, and then analyzed the relationships using linear regression analysis.

### Statistical analysis

GraphPad Prism software was used for all statistical analyses. All measurements were taken from distinct samples unless otherwise stated. All data were tested for normality.Pairwise analyses used two-tailed paired t-tests (animal level data, baseline vs. treatment) or Mann-Whitney tests (pooled cell level data, baseline vs. treatment). Group analysis data used two-way ANOVA with Tukey’s multiple comparisons tests (correlations between different cell types within group), one-way ANOVA with Dunnett’s multiple comparisons tests (cell level data correcting for multiple comparisons), Fisher’s LSD tests (animal level data not correcting for multiple comparisons) (comparing different drug treatment groups to the control group), or one-way repeated measures ANOVA with Tukey’s multiple comparisons tests (comparing multiple treatments within the same group correcting for multiple comparisons). All regression analysis was performed using simple linear regression. Measures were reported as mean ± standard error of the mean, unless otherwise indicated.

## Acknowledgements

We thank M.P. Cousineau and S. Baillet (McGill University) for assistance with Brainstorm software. Artwork was kindly provided by Ms. Audrey Desaulniers, Orcéine, Montreal, Canada. This research was supported by Canada Graduate Scholarships-Masters to N.B. and V.L. and a Postgraduate Scholarship - Doctoral to N.B from the Natural Sciences and Engineering Research Council of Canada. E.S.R. is funded by a grant from the Canadian Institutes of Health Research (FDN-143238) and a Research Chair from the Fonds de recherche santé Québec (31036). A.S. and E.S.R. were also supported by the Brain Canada Canadian Neurophotonics Platform.

## Contributions

N.B. and E.S.R. conceived and designed the study. N.B. performed all imaging experiments and data analysis. A.S. developed the labeling methodology and provided technical support. V.L. provided analysis software and helped develop labeling methodology. N.B. and E.S.R. drafted the manuscript. All authors provided critical feedback and editing of the final manuscript. E.S.R. supervised the research.

